# A conundrum of pleiotropy: GWAS-derived pleiotropy is weakly negatively associated with co-expression network centrality in *Arabidopsis thaliana*

**DOI:** 10.64898/2026.07.15.738774

**Authors:** Chatendeep Gill, Samuel Yeaman

## Abstract

Pleiotropy, the influence of a single gene on multiple phenotypic traits, is a fundamental feature of genetic systems but remains difficult to quantify at genome-wide scales. Genome-wide association studies (GWAS) provide one avenue to characterize pleiotropy through multi-trait genetic associations, while functional genomic approaches such as gene co-expression networks offer indirect predictions of gene importance and pleiotropic potential based on network centrality statistics. However, the relationship between GWAS-derived pleiotropy and network-based measures of gene importance remains unclear. Here, we quantified gene-level pleiotropy in *Arabidopsis thaliana* using gene-trait associations from the AraGWAS Catalog and AraPheno databases and compared them to previous estimates of pleiotropy based on gene co-expression networks. GWAS pleiotropy was measured using the Hill number of order *q* = 1, an entropy-based metric adapted from ecological diversity theory. Across 2,179 genes, pleiotropy exhibited a right-skewed distribution, with most genes influencing relatively few traits and a smaller subset exhibiting broad multi-trait effects. Contrary to expectations, pleiotropy was weakly negatively correlated with multiple measures of co-expression network centrality, including degree, strength, closeness, and betweenness. Additional analyses showed that GWAS-associated genes were enriched within the co-expression network, exhibited broader expression across tissues, and were modestly enriched among conserved BUSCO genes. Together, these findings reveal a conundrum: although genes identified by GWAS are broadly expressed and integrated into functional networks, the most pleiotropic genes occupy relatively peripheral positions within those networks. This unexpected pattern suggests that GWAS-derived pleiotropy captures a distinct aspect of gene function than co-expression network centrality.

## Introduction

Pleiotropy, the phenomenon in which a single gene influences multiple phenotypic traits, is a fundamental property of genetic systems and plays an important role in shaping evolutionary dynamics (Paaby & Rockman, 2013; Pavlicev & Wagner, 2012). As genes participate in interconnected biological processes, mutations often have effects that extend across multiple traits, generating genetic correlations that can constrain or facilitate evolutionary responses to selection (Wessinger & Hileman, 2016; Zhang, 2023). Consequently, the distribution of pleiotropic effects across genes has important implications for evolutionary theory, influencing the rate of adaptation and shaping the genetic architecture of adaptation (Hansen, 2003) as well the “cost of complexity,” which predicts that mutations affecting many traits experience stronger selective constraints than mutations with more limited effects (Orr, 2000; Orr, 2005). Evaluating these theoretical predictions requires reliable estimates of pleiotropy across large numbers of genes.

Despite its importance, estimating pleiotropy at genome-wide scales remains challenging. One widely used approach is to infer pleiotropy from genome-wide association studies (GWAS), in which genes associated with multiple phenotypic traits are interpreted as having broader pleiotropic effects. Large-scale GWAS have revealed that pleiotropy is pervasive across complex traits, identifying hundreds of loci associated with multiple phenotypes and enabling the characterization of shared genetic architecture, causal relationships, and genetic overlap among traits (Gratten & Visscher, 2016; Schaid et al., 2016; Visscher & Yang, 2016; Zhu et al., 2021). An alternative approach estimates pleiotropy indirectly from functional genomic data. Gene co-expression networks use patterns of coordinated gene expression to infer functional connectivity, with highly connected “hub” genes predicted to participate in numerous biological processes and therefore exhibit greater pleiotropy (Mähler et al., 2017; Muzio et al., 2023). These complementary approaches have increasingly been applied to address evolutionary questions, including how pleiotropy shapes adaptation and parallel evolution (Lai et al., 2025). However, it remains unclear whether GWAS-derived and network-based estimates quantify the same underlying biological phenomenon or instead capture different dimensions of gene function.

Here, we compare gene-level estimates of pleiotropy obtained from these two complementary approaches. Using *Arabidopsis thaliana* as a model system, we quantified pleiotropy from gene-trait associations in the AraGWAS Catalog (Togninalli et al., 2017) and AraPheno (Togninalli et al., 2019) and compared these estimates with measures of gene co-expression network centrality. Simply counting the number of associated traits may overestimate pleiotropy when a gene has one or a few strong associations but many additional weak associations that are unlikely to reflect broad biological influence. To account for this, we quantified GWAS-derived pleiotropy using the Hill number of order *q* = 1, an entropy-based metric adapted from ecological diversity theory that incorporates both trait richness and the distribution of genetic effects among traits (Alberdi & Gilbert, 2019; Chao et al., 2014; Jost, 2006). This framework distinguishes genes whose effects are distributed relatively evenly across many traits from those whose apparent pleiotropy is driven by one or a few strong associations accompanied by numerous weak associations.

The primary objective of this study was to determine whether GWAS-derived estimates of pleiotropy correspond to network-based estimates of gene pleiotropy. Specifically, we tested whether highly pleiotropic genes occupy central positions within gene co-expression networks. To provide additional context for patterns of GWAS detectability, we also examined relationships between GWAS representation, tissue expression breadth, and evolutionary conservation. Together, these analyses provide a genome-wide comparison of two widely used approaches for estimating pleiotropy and evaluate whether they identify the same genes as functionally important.

## Materials and Methods

### Data sources

Genome-wide association data were obtained from the AraGWAS Catalog, a curated repository of SNP-trait associations in *Arabidopsis thaliana* (Togninalli et al., 2017, 2019). Trait metadata, including trait descriptions and classifications, were obtained from the AraPheno database, which provides standardized phenotypic information linked to AraGWAS entries. Gene co-expression network statistics and tissue specificity estimates (τ) were obtained from Nocchi et al. (2024), who calculated these metrics using Arabidopsis gene expression data from previously published datasets (Papatheodorou et al., 2018). Co-expression metrics included node degree, node strength, closeness centrality, and betweenness centrality. Tissue specificity (τ) was used as a measure of expression breadth, where lower values indicate broader expression across tissues and higher values indicate greater tissue specificity. A genome-wide background set of protein-coding genes was obtained from the TAIR10 *Arabidopsis thaliana* protein annotation available through Ensembl Plants (Cheng et al., 2017; Yates et al., 2026). Gene identifiers were extracted from all annotated protein-coding loci and used as the genomic background for enrichment analyses involving co-expression network genes and BUSCO genes. To evaluate whether highly conserved genes were disproportionately represented among AraGWAS genes, Benchmarking Universal Single-Copy Orthologs (BUSCOs) were identified from the embryophyta lineage dataset (Manni et al., 2021; Tegenfeldt et al., 2025). Genes classified as Complete or Duplicated BUSCOs were retained. Transcript identifiers were converted to TAIR10 gene identifiers by removing transcript-specific suffixes, and duplicate entries were removed prior to analysis.

### Estimating effect size per gene and per trait

To estimate pleiotropy for each gene, it was necessary to build a *gene* × *trait* matrix in which each entry represented the effect size of a given gene on a given trait. To populate this matrix, SNP-trait associations were filtered to retain statistically significant signals using a threshold of *p* < 1 × 10^-4^, consistent with the definition of high-confidence associations in the AraGWAS Catalog. Where multiple SNPs with significant associations for a given trait mapped to the same gene, these effects were aggregated by calculating the mean absolute standardized effect size (mean |*z*|). This approach assumes that when there are multiple significant GWAS hits for the same trait within a given gene, they most likely arise from linkage to a causal variant. GWAS datasets frequently include measurements of multiple closely related phenotypes across different timepoints or experimental conditions (*e*.*g*., root length at timepoints x, y, z). Treating these measurements as independent traits could inflate estimates of pleiotropy by increasing the apparent number of traits associated with a gene. To address this, measurements sharing the same trait ontology annotation in AraPheno were collapsed by calculating the mean of the mean absolute standardized effect size (mean (mean |*z*|)), which was used as the final effect size for each gene-trait combination.

### Quantifying pleiotropy using Hill numbers

Gene-level pleiotropy was quantified using the Hill number of order *q* = 1, equivalent to the exponential of Shannon entropy (Chao et al., 2014; Jost, 2006). For each gene, Shannon entropy (*H*) was calculated from the normalized distribution of gene-trait effect sizes:

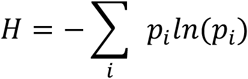

where *p*_*i*_ represents the proportion of the gene’s total effect attributed to trait *i*. The Hill number (*q* = 1) was then calculated as:

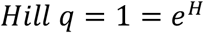

This metric estimates the effective number of traits influenced by a gene incorporating both trait richness (the number of traits with non-zero effects) and evenness (the distribution of effect sizes across these traits). Genes with no detectable effects across all traits (total effect = 0) were excluded, as entropy-based metrics require a defined probability distribution.

All analyses were performed in R (R Core Team, 2025).

## Results and Discussion

Gene-level GWAS pleiotropy, quantified using the Hill number of order *q* = 1, varied widely across the genome. The distribution of pleiotropy was slightly right-skewed, with most genes exhibiting low to moderate pleiotropy and a smaller subset showing substantially higher values (Figure 1A). Across the 2,179 genes analyzed, the mean Hill number was 9.26 and the median was 8.97. The 5 most pleiotropic genes exhibited a Hill number of 18.79 - 20.62 (Table 1). This pattern suggests that the majority of genes influence relatively few traits, whereas a minority contribute to broad, multi-trait effects. Highly pleiotropic genes were associated with diverse trait categories, including flowering time, root morphology, growth and development, ionomics, and stress or disease resistance.

**Table 1.** Top five most pleiotropic genes identified in this study, ranked by Hill number (*q* = 1). Listed traits represent broad categories of associated phenotypes, illustrating the diversity of biological processes influenced by highly pleiotropic genes.

| Gene | Hill $q = 1$ | Representative Trait Categories |
| --- | --- | --- |
| AT1G32170 | 20.62 | Flowering time<br>Root traits<br>Growth and development<br>Ionomics<br>Stress/Disease resistance |
| AT3G44400 | 19.67 | Flowering time<br>Root traits<br>Growth and development<br>Ionomics<br>Leaf traits |
| AT5G45100 | 18.91 | Flowering time<br>Root traits<br>Ionomics<br>Stress/Disease resistance<br>Seed dormancy/germination |
| AT5G46270 | 18.89 | Flowering time<br>Root traits<br>Growth and development<br>Ionomics<br>Stress/Disease resistance<br>Seed dormancy/germination<br>Leaf traits |
| AT3G44620 | 18.79 | Flowering time<br>Root traits<br>Growth and development<br>Ionomics<br>Stress/Disease resistance<br>Seed dormancy/germination<br>Leaf traits |

**Figure 1.**
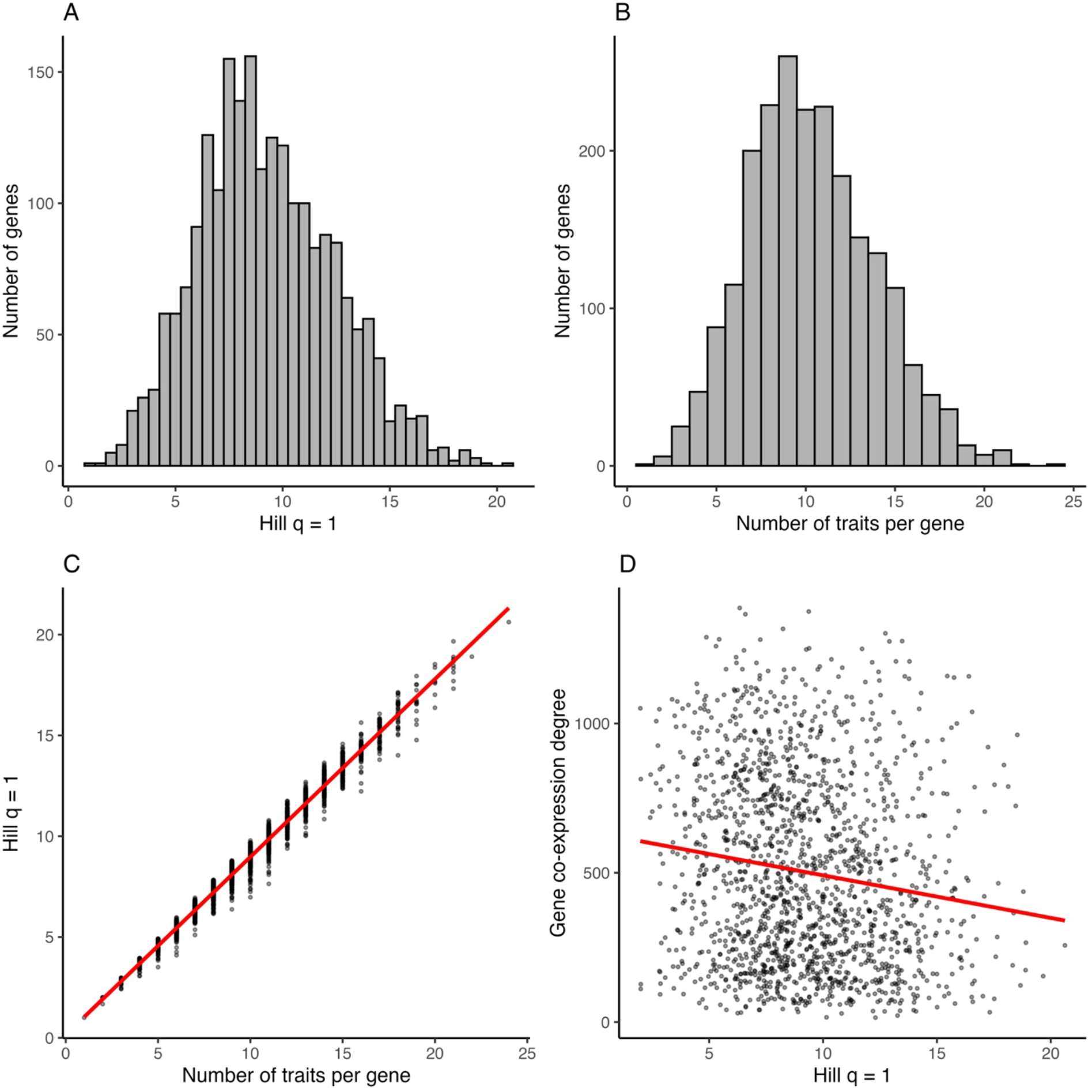
Genome-wide distribution and determinants of gene pleiotropy in *Arabidopsis thaliana*. (A) Distribution of gene-level pleiotropy quantified using the Hill number (*q* = 1), showing a slightly right-skewed distribution in which most genes exhibit low to moderate pleiotropy and a small subset exhibit high pleiotropy. (B) Distribution of the number of traits associated with each gene following trait aggregation. (C) Relationship between trait richness and Hill number (*q* = 1), demonstrating a strong, near-linear relationship between the number of associated traits and estimated pleiotropy. (D) Relationship between Hill number (*q* = 1) and gene co-expression network centrality (node degree), showing that genes with higher pleiotropy tend to occupy less connected positions within the co-expression network. **ALT TEXT:** Four-panel figure comparing gene pleiotropy, trait richness, and network connectivity. Panels A and B show histograms with unimodal, slight right-skewed distributions. Panel C shows a scatterplot with points closely aligned along an upward-sloping fitted line. Panel D shows a scatterplot with widely dispersed points and a downward-sloping fitted line.

Given the expectation that highly pleiotropic genes occupy central positions within biological networks, we compared GWAS-derived pleiotropy with measures of gene co-expression network centrality. Unexpectedly, GWAS-derived pleiotropy exhibited weak but statistically significant negative correlations with multiple measures of network centrality, including node degree (Figure 1D; Spearman’s *ρ* ≈ -0.152, *p* = 6.02 × 10^-10^), node strength (Spearman’s *ρ* ≈ -0.154, *p* = 3.15 × 10^-10^), closeness (Spearman’s *ρ* ≈ -0.108, *p* = 1.10 × 10^-5^), and betweenness (Spearman’s *ρ* ≈ -0.051, *p* = 0.039). Although modest in magnitude, these consistent negative relationships indicate that genes influencing many traits tend to occupy slightly less connected positions within the co-expression network. This result contrasts with network-based models in which highly connected genes are expected to participate in a broader range of biological processes and therefore exhibit greater pleiotropy (Mähler et al., 2017). However, it is broadly consistent with previous analyses showing that GWAS-associated genes occupy relatively peripheral positions within biological interaction networks (Barrenas et al., 2009).

To determine whether this pattern depended on how pleiotropy was quantified, we examined its relationships with trait richness and effect size. As expected, the Hill number was strongly associated with the number of traits linked to each gene (Figure 1B and 1C; Spearman’s *ρ* ≈ 0.99, *p* < 2.2 × 10^-16^), demonstrating that genes associated with more traits generally exhibited greater pleiotropy. This result is consistent with ecological applications of Hill numbers, where richness often dominates diversity estimates when abundance distributions are highly uneven (Chao et al., 2014; Jost, 2006). Consequently, GWAS-derived pleiotropy should be interpreted primarily as a measure of the breadth of traits influenced by a gene rather than the evenness of its effects across traits.

Likewise, total effect magnitude increased with pleiotropy (Spearman’s *ρ* ≈ 0.65, *p* < 2.2 × 10^-16^), reflecting the cumulative effects of genes influencing larger numbers of traits. In contrast, after standardizing effect size by the number of associated traits, mean effect size per trait showed no meaningful relationship with pleiotropy (Spearman’s *ρ* ≈ -0.04, *p* = 0.09), indicating that highly pleiotropic genes do not exert stronger effects on individual traits. Instead, their greater cumulative effects arise primarily from influencing more traits rather than from stronger per-trait effects. Mean effect size per trait also showed only a weak positive relationship with co-expression network degree (Spearman’s *ρ* ≈ 0.07, *p* = 0.005), indicating that genes occupying more central network positions do not generally exert substantially stronger effects on individual traits. Moreover, the weak negative relationship between co-expression network degree was also observed when trait richness, rather than the Hill number, was used as the measure of pleiotropy (Spearman’s *ρ* ≈ -0.16, *p* = 1.0 × 10^-10^), indicating that the negative association with network centrality was robust to alternative measures of gene-level pleiotropy. Together, these findings suggest that highly pleiotropic genes primarily influence many traits through numerous modest effects rather than a small number of exceptionally large ones, consistent with previous empirical work (Paaby & Rockman, 2013; Wang et al., 2010; Zhang, 2023).

The unexpected negative relationship between GWAS-derived pleiotropy and network centrality could have arisen if genes under the strongest purifying selection were underrepresented in GWAS datasets, consistent with previous suggestions that the relative peripherality of GWAS genes reflects stronger selective constraints on highly central genes (Barrenas et al., 2009). To explore this possibility, we first examined whether genes represented in the co-expression network were more likely to be detected by GWAS. Of the 18,571 genes represented in the co-expression network, 1,646 (8.9%) were associated with at least one AraGWAS trait. In contrast, only 523 of 9,057 genes absent from the co-expression network (5.8%) were associated with a GWAS hit. Co-expression network genes were therefore significantly enriched for GWAS-associated genes (*odds ratio* = 1.59, 95% *CI* = 1.43 – 1.76, *p* < 2.2 × 10^-16^), with the network containing 1,646 of 2,169 (75.9%) of all GWAS-associated genes.

We next examined whether tissue expression breadth influenced GWAS detectability. Genes with at least one GWAS association exhibited lower mean tissue specificity (τ = 0.505) than genes without GWAS associations (τ = 0.571), indicating that broadly expressed genes were more likely to be detected by GWAS. Logistic regression supported this pattern, with tissue specificity showing a significant negative association with GWAS detectability (*β* = -0.774 ± 0.080 *SE, z* = -9.65, *p* < 2 × 10^-16^). A one-unit increase in τ reduced the odds of GWAS detection (*odds ratio* = 0.46, 95% *CI* = 0.39 – 0.54).

Finally, we tested whether highly conserved genes were disproportionately represented among GWAS-associated genes using BUSCO annotations. Of the 1,630 complete or duplicated BUSCO genes, 156 (9.6%) were represented in AraGWAS. In contrast, 7.7% of non-BUSCO genes were represented in AraGWAS. BUSCO genes were therefore modestly but significantly enriched among GWAS-associated genes (*odds ratio* = 1.26, 95% *CI* = 1.06 – 1.50, *p* = 0.009). Together, these analyses indicate that although GWAS preferentially detects broadly expressed and evolutionarily conserved genes, these biases are unlikely to explain the unexpected negative relationship between GWAS-derived pleiotropy and co-expression network centrality.

Therefore, the negative relationship between pleiotropy and network centrality cannot be explained simply by the underrepresentation of constrained genes in GWAS datasets (Mähler et al., 2017; Barbitoff et al., 2025). Instead, our results extend previous observations that GWAS-associated genes occupy relatively peripheral network positions (Barrenas et al., 2009) by demonstrating that this pattern persists even among genes with the highest levels of GWAS-derived pleiotropy. Although both approaches are widely used to estimate gene pleiotropy, they appear to quantify complementary rather than equivalent aspects of gene function. More broadly, these results challenge the assumption that different estimates of pleiotropy necessarily capture the same underlying biological phenomenon (Paaby & Rockman, 2013; Zhang, 2023) and emphasize the importance of considering multiple approaches when investigating the genetic architecture of complex traits.

Several limitations should be considered when interpreting these findings. GWAS-based analyses are subject to internal constraints including limited statistical power, linkage disequilibrium, and confounding among correlated traits (Reinert, 2022; Schaid et al., 2016). Although trait aggregation reduced inflation of pleiotropy estimates arising from repeated measurements of the same biological trait, this methodological choice may also obscure biologically meaningful differences in gene effects across different developmental stages, environmental conditions, or other experimental contexts. In addition, the present framework does not explicitly account for trait correlations, as implemented in alternative approaches such as HOPS (Jordan et al., 2019). Consequently, some degree of statistical pleiotropy may remain in the dataset. These limitations should be considered when interpreting the magnitude of the observed relationships but are unlikely to alter the overall conclusion that GWAS-derived pleiotropy and network centrality capture distinct aspects of gene function.

Future research should determine whether the disconnect observed here between GWAS-derived pleiotropy and network centrality is a general feature of genetic architecture or is specific to *Arabidopsis thaliana* and the datasets analyzed. Applying comparable analyses across additional species, trait datasets, and network resources would help determine the extent to which network position predicts observable pleiotropic variation. More broadly, integrating GWAS, expression, and network data may provide a useful framework for distinguishing genes that are functionally central from those that contribute disproportionately to standing phenotypic variation.

## Funding

This research was supported by a Natural Sciences and Engineering Research Council of Canada (NSERC) Undergraduate Student Research Award (USRA) awarded to Chatendeep Gill and by an NSERC Discovery Grant (RGPIN-03310-2023) awarded to Samuel Yeaman.

## Conflict of Interest

The authors declare no conflicts of interest.

## Data Availability

The data underlying this article are available in GitHub at https://github.com/ChatendeepGill/arabidopsis-gwas-pleiotropy. Original AraGWAS Catalog data are not redistributed in this repository and are available from the AraGWAS Catalog (https://aragwas.1001genomes.org/).

## Notes

### Competing Interest Statement

The authors have declared no competing interest.

